# SurpHer: a genetically encoded ratiometric sensor for dynamic extracellular pH imaging

**DOI:** 10.64898/2026.05.18.725923

**Authors:** Sofie Cens Holste, Leïla Dos Santos, Mahdi Rezayati Charan, Oline Nyhegn-Eriksen, Roxane Crouigneau, Birthe B. Kragelund, Rodolphe Marie, Albin Sandelin, Jamie Yam Auxillos, Stine Falsig Pedersen

## Abstract

Extracellular pH is a key microenvironmental factor shaping cell physiology and disease, creating a need for quantitative biosensors that can capture dynamic changes in pH_e_ at the surface of individual living cells. Here, we develop a genetically encoded, ratiometric extracellular pH biosensor through systematic screening of a modular library of membrane-display designs that combine SEpHluorin with a pH-stable reference fluorophore. Screening identified a cell-surface-localised mKate2-SEpHluorin construct, named SurpHer, that exhibits dynamic ratiometric responses across the pH_e_ range of 6 - 7.8. SurpHer shows robust membrane localisation and extracellular pH responsiveness across diverse human cell types including HEK293T, PANC-1 and MDA-MB231 cells. Following stable integration in MDA-MB-231 cells, SurpHer enabled time-course imaging of pH_e_ gradients in a microfluidic platform for modelling tumour microenvironments. SurpHer enables real-time interrogation of the pericellular pH environment of tumor cells and, more broadly, provides a strategy to probe microenvironmental pH dynamics across diverse biological contexts.

## Introduction

Extracellular pH (pH_e_) is a key regulator of cellular behaviour across diverse physiological and pathological microenvironments. In mammalian tissues, pH_e_ is typically maintained within a narrow range of approximately 7.3-7.4. However, local pH_e_ deviations arise in normal physiology, for example, in organs engaged in extensive acid–base transport, such as the stomach and kidney, and in skeletal muscle during exercise (Swietach, Boedtkjer and Pedersen, 2023). In disease, pH_e_ dysregulation is associated with inflammatory microenvironments and is particularly pronounced in solid tumours, where increased metabolic acid production and poor perfusion drive extracellular acidification. This acidic microenvironment influences tumour invasion, extracellular matrix remodelling, immune cell function and therapeutic response (Corbet *et al*., 2020; Czaplinska *et al*., 2023; Swietach, Boedtkjer and Pedersen, 2023; Stigliani *et al*., 2024). Together, these highlight pH_e_ as a biologically and clinically important microenvironmental variable, whose spatiotemporal dynamics remain insufficiently resolved.

Despite its broad biological relevance, visualising pH_e_ dynamics at single-cell resolution in real time remains challenging. Microelectrodes (Jensen *et al*., 1997; Yang *et al*., 2021) and whole-tissue imaging modalities such as magnetic resonance imaging and positron emission tomography (Anemone *et al*., 2019; Crouigneau *et al*., 2024) have provided important insights into extracellular pH regulation, but are invasive or poorly suited to longitudinal single-cell measurements. Cell surface-adhering fluorescent pH sensitive dyes (Ke *et al*., 2014; Yang *et al*., 2018) exist which enable ratiometric quantification of pericellular pH at single-cell resolution, but these are best suited to relatively simple systems or short-duration experiments and may perturb the cells they are intended to monitor.

Genetically encoded reporters offer an attractive alternative because they can be expressed directly by cells and maintained over the course of longitudinal experiments. However, existing pH_e_ biosensor designs still impose important trade-offs. Single-fluorophore pH_e_ biosensors targeted to the plasma membrane are limited by their non-ratiometric readout, which causes the fluorescence readout to be affected by variation in sensor abundance, cell density and optical conditions (Shen *et al*., 2014), limiting their use to short-term, homogenous two-dimensional cell culture studies. FRET-based sensors provide a ratiometric alternative, but commonly used FRET pairs can introduce additional sensitivities; for example, enhanced yellow fluorescent protein (EYFP) fluorescence is influenced by both pH and chloride (Kuner and Augustine, 2000). EGF-fused pH sensor was recently reported but requires high EGFR expression and incubation with recombinantly produced EGF fusion protein (Karsten *et al*., 2022), potentially affecting cell phenotypes. Together, these trade-offs leave a need for a genetically encoded pH_e_ biosensor.

Here, we report SurpHer, a genetically encoded, membrane-anchored ratiometric pH biosensor for quantitative pericellular pH imaging. Identified from systematic screening of a modular library of plasma-membrane-targeted biosensor designs, SurpHer localises robustly to the cell surface and responds across the physiologically relevant extracellular pH range. We show that SurpHer performs across multiple cell types and enables real-time longitudinal imaging of pH_e_ gradients in a microfluidic tumour-microenvironment model. SurpHer, therefore, is a versatile tool for resolving pH_e_ dynamics in complex biological systems.

## Results

### Optimising a ratiometric pH_e_ biosensor for plasma membrane localisation

To develop a genetically encoded biosensor for reporting pH_e_ at the cell surface, we designed a modular library of plasma membrane-targeted ratiometric constructs in which the pH-sensitive fluorophore SEpHluorin was paired with a reference fluorophore and combined with distinct leader, linker and anchor sequences to promote extracellular display at the plasma membrane (Fig. 1A,B). We varied promoter strength (pEF1a or pCMV promoters), reference fluorophore type (mCherry or mKate2), linker composition (serine-glycine linker or linker218 (Whitlow *et al*., 1993)), and membrane-targeting strategy, combining IgK, CD59 or HA leader sequences with the transmembrane domains of the platelet-derived growth factor receptor (PDGFR) or the glycosylphosphatidylinositol (GPI) anchors of the cell death receptor CD59 (CD59-GPI), or the reticulon-4 receptor (Nogo receptor, NGR), respectively (Fig. 1B). The CD59 leader/CD59-GPI and HA leader/NGR-GPI targeting combinations (Nasu *et al*., 2021), together with linker218 (Whitlow *et al*., 1993), were included based on prior reports supporting efficient membrane display of cell-surface proteins. The generated constructs were first screened in HEK293T cells by live-cell epifluorescence microscopy to assess co-localisation of the pH-sensitive and reference fluorophores at the plasma membrane.

**Figure 1:**
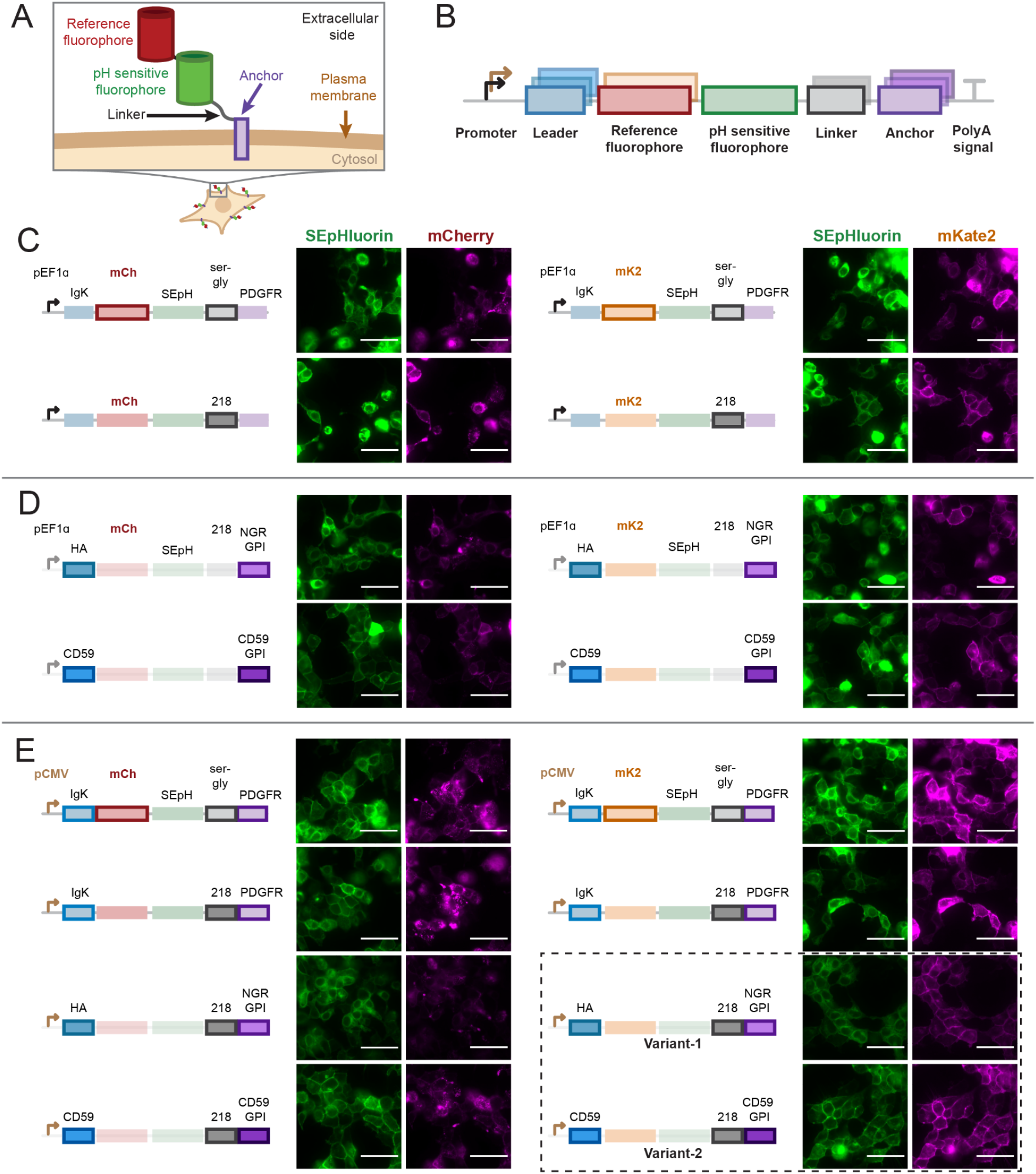
Design and imaging-based screening of a plasma membrane-targeted extracellular pH biosensor library. **A**. Schematic of the membrane-anchored biosensor design, showing extracellular display of the pH-sensitive fluorophore, reference fluorophore, linker and membrane anchor at the plasma membrane. **B**. Modular architecture of the construct library. Promoter, leader sequence, reference fluorophore identity, linker and membrane anchor were varied. **C**. Representative live-cell fluorescence images of pEF1a/IgK leader/PDGFR anchor constructs comparing reference fluorophore identity and linker composition. Left, mCherry-based constructs; right, mKate2-based constructs. **D**. Representative live-cell fluorescence images of pEF1a linker218 constructs with alternative leader-anchor combinations. **E**. Representative live-cell fluorescence images of pCMV-driven constructs comparing fluorophore identity, linker composition and leader-anchor combinations. Dashed box indicates shortlisted candidates, variant 1 and variant 2. Scale bars, 50 μm. Imaged with 40x objective. Images were contrast-adjusted for display using channel-specific minimum and maximum intensity settings.

Initial optimisation focused on reference fluorophore identity and linker composition with a fixed pEF1a/IgK leader/PDGFR anchor architecture. In the mCherry-based constructs, intracellular aggregates were observed with both linkers, resulting in poor fluorescence overlap between channels (Fig. 1C, left). By contrast, mKate2-based constructs showed improved plasma membrane-associated fluorescence, particularly when paired with linker218, however this was not consistently observed across all cells (Fig. 1C, right). We therefore prioritised linker218 for subsequent optimisation.

We then tested whether alternative leader-anchor combinations improved cell-surface localisation with the linker218. Under the pEF1a promoter, both the HA/NGR-GPI and CD59/CD59-GPI combinations showed membrane-associated fluorescence across both channels, primarily in the mKate2-based designs, although intracellular localisation remained evident in a subset of cells (Fig. 1D).

Finally, we asked whether promoter choice further influenced biosensor localisation. When the same biosensor architectures were placed under the control of the pCMV promoter, mCherry-based constructs still showed intracellular puncta, whereas the corresponding mKate2-based constructs displayed strong plasma membrane localisation with reduced intracellular aggregates (Fig. 1E). Among these, the pCMV-HA leader-mKate2-SEpHluorin-linker218-NGR GPI anchor construct (**variant 1**) and the pCMV-CD59 leader-mKate2-SEpHluorin-linker218-CD59 anchor construct (**variant 2**) emerged as the strongest candidates, with primarily plasma membrane localisation of both fluorophores in the majority of cells and minimal intracellular aggregates or diffuse intracellular fluorescence (Fig. 1E, dotted outline). These variants were therefore selected for further characterisation.

### Biosensor variants 1 and 2 exhibit plasma membrane localisation across different cell lines

To assess biosensor performance and generalisability across cell types, we characterised the two candidates, (variant 1 and variant 2), after transiently transfecting into HEK293T, PANC-1 and MDA-MB231 cells. This allowed us to evaluate whether the localisation behaviour observed during the initial screen was retained across distinct cellular contexts, including cell lines relevant for modelling pH_e_ dysregulation in cancer. Across all three cell types and both variants, live-cell images showed predominantly peripheral fluorescence in the SEpHluorin and mKate2 channels, although intracellular signal was evident in some cells (Fig. 2A; Supplementary Figs. 1A–5A). Line-profile analysis further confirmed enrichment of both fluorophores at the cell boundary across the validation set, supporting plasma membrane-associated localisation of the ratiometric reporter (Fig. 2A; Supplementary Figs. 1A–5A).

**Figure 2:**
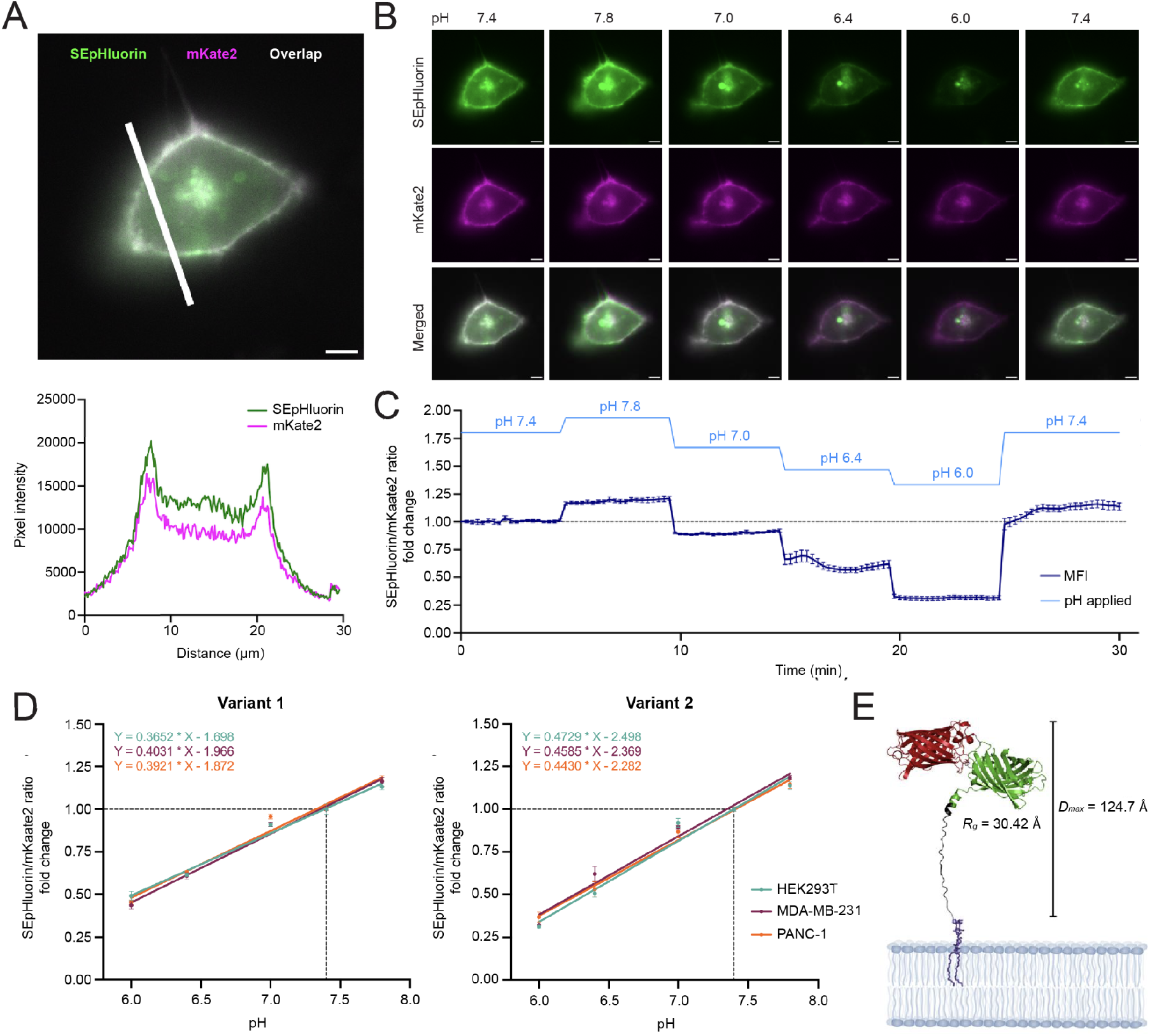
SurpHer pH_e_-sensitive genetically encoded ratiometric biosensor validation in 2D MDA-MB-231 cells. **A**. Representative validation of membrane localization of the biosensor. SEpHluorin in green, mKate2 in magenta and overlap in white. Scale bar: 5 µm. The white line represents the ROI for the profile plot analysis. **B**. Representative images of transiently transfected MDA-MB-231 cells in pH 7.8, 7.4, 7.0, 6.4 and 6.0. Imaged with a 60x objective. pH insensitive fluorophore: mKate2 (ex: 588 nm, em: 633 nm). pH sensitive fluorophore: SEpHluorin (ex: 488 nm, em: 511 nm). Scale bar: 5 µm. **C**. Quantification of pHe-sensitivity during exposure to pH 7.8, 7.4, 7.0, 6.4 and 6.0. Images acquired every 15 sec for a total of 30 min. Data analysis by ROI marking of the cell membrane for intensity measurement during pH exposures over time, MFI = mean fluorescence intensity. Normalization to first measurement in pH 7.4. n = 3, SEM error bars. **D**. Linear regression of normalized SEpHluorin/mKate2 ratio versus extracellular pH for sensor variants 1 (left) and variant 2 (right) in HEK293T, MDA-MB-231 and PANC-1 cells exposed to Ringer solutions at pH 7.8, 7.4, 7.0, 6.4 and 6.0. Both variants showed strong linear pH-dependent responses across all cell lines (R^2^ = 0.965–0.983). Slopes did not differ significantly between cell lines within variant 1 (p = 0.2306) or variant 2 (p = 0.6046), whereas variant 2 exhibited significantly steeper slopes than variant 1 across cell lines (p = 0.0004). Variant 2 slopes ranged from 0.443–0.473 compared with 0.365–0.403 for variant 1. Simple linear regression. n = 3, SD error bars. **E**. Visualization of the GPI-anchored extracellular biosensor domain. The AlphaFold 3-predicted protein model was combined with a separately modelled GPI anchor (default CHARMM-GUI core sequence: PI–GlcN–3×Man–EtNP), shown in the context of a schematic representation of the plasma membrane. Note that the position of the protein relative to the membrane was set manually and is illustrative; the actual conformational ensemble is not represented. CRYSOL analysis of the extracellular biosensor domain only (excluding the GPI anchor) yielded Rg=30.4 Å and a maximum molecular dimension of 124.7 Å.

### SurpHer exhibits dynamic pH_e_ sensitivity and reversibility across different cell lines

To assess pH responsiveness, we performed live-cell imaging in Ringer solutions spanning pH 6.0–7.8, covering physiologically and pathologically relevant pH_e_ conditions. For both variants, the SEpHluorin/mKate2 ratio decreased stepwise with decreasing pH_e_ and rebounded upon return to neutral conditions, demonstrating reversible pH_e_ reporting at the cell surface (Fig. 2B-C; Supplementary Figs. 1B-5B). Notably, the ratio shifted detectably within seconds after each Ringer solution exchange, indicating that the biosensor rapidly reports dynamic pH_e_ changes under live-cell imaging conditions (Fig. 2C; Supplementary Figs. 1C-5C). All cell lines (HEK293T, MDA-MB-231, and PANC-1) exhibited strong linear relationships between pH and the normalized SEpHluorin/mKate2 ratio for both sensor variants (R^2^ = 0.965–0.983) (Fig. 2D). No significant differences in slope were observed between cell lines within either variant (variant 1: p = 0.2306; variant 2: p = 0.6046), indicating no cell line–dependent variation in pH responsiveness. In contrast, comparison between variants showed a significant difference in slope (p = 0.0004), demonstrating distinct pH sensitivities between variant 1 and variant 2. Variant 2 consistently exhibited higher slopes across all three cell lines (0.443–0.473) compared to variant 1 (0.365–0.403), indicating a greater change in the normalized SEpHluorin/mKate2 ratio per unit change in pH (Fig. 2D). Together, these results demonstrate that variant 2 confers an increased pH-dependent response magnitude that is consistent across cellular contexts (Figure 2D). Variant 2 was therefore selected for further characterisation and hereafter termed SurpHer.

To provide a structural context for the selected SurpHer design, we generated a series of AlphaFold 3 (AF3) models of the extracellular biosensor domain and from selection based on highest pTM confidence scores, we assembled a visualization of the GPI-anchored form of SurpHer by combining the AF3 model with a separately generated GPI anchor. CRYSOL analysis of the extracellular SurpHer domain, excluding the GPI anchor and bilayer, yielded a radius of gyration (Rg) of 30.4 Å and a maximum molecular dimension of 124.7 Å (Figure 2E).

We next assessed the pH sensitivity and imaging stability of the reference fluorophore, as ratiometric measurements depend on the behaviour of both fluorescent channels. mKate2 fluorescence exhibited minor pH-dependent variation, consistent with previous reports (Shcherbo *et al*., 2009), although substantially smaller than the SEpHluorin fluorescence change across the tested extracellular pH range (Supplementary Figs. 6–7). We then assessed the photostability of SurpHer under repeated live-cell imaging. Under standard imaging conditions, the mean membrane fluorescence intensity of the SEpHluorin channel remained largely stable over 15 min and the mKate2 channel showed only a slight decrease in intensity across HEK293T, MDA-MB-231 and PANC-1 cells, supporting its use for longitudinal live-cell imaging over this timescale (Supplementary Fig. 8A–C). Under higher-intensity illumination, mean fluorescence intensity declined more substantially, particularly in the mKate2 channel (Supplementary Fig. 8D–F).

### Stable expression of SurpHer enables extracellular pH imaging in a microfluidic platform mimicking tumour microenvironment pH gradients

One advantage of a genetically encoded pH_e_ sensor is its ability to support longitudinal imaging. To establish a stable reporter cell line suitable for longer-term studies in microfluidic systems, we cloned the SurpHer biosensor, together with a puromycin resistance cassette, into a piggyBac expression vector for genomic integration in MDA-MB-231 cells (Materials and Methods, Fig. 3A). Following nucleofection with both the SurpHer piggyBac expression vector and the transposase vector, puromycin selection and fluorescence-activated cell sorting (FACS) were used to select cells expressing both SEpHluorin and mKate2 (Figs. 3A,B) at a high level.

**Figure 3:**
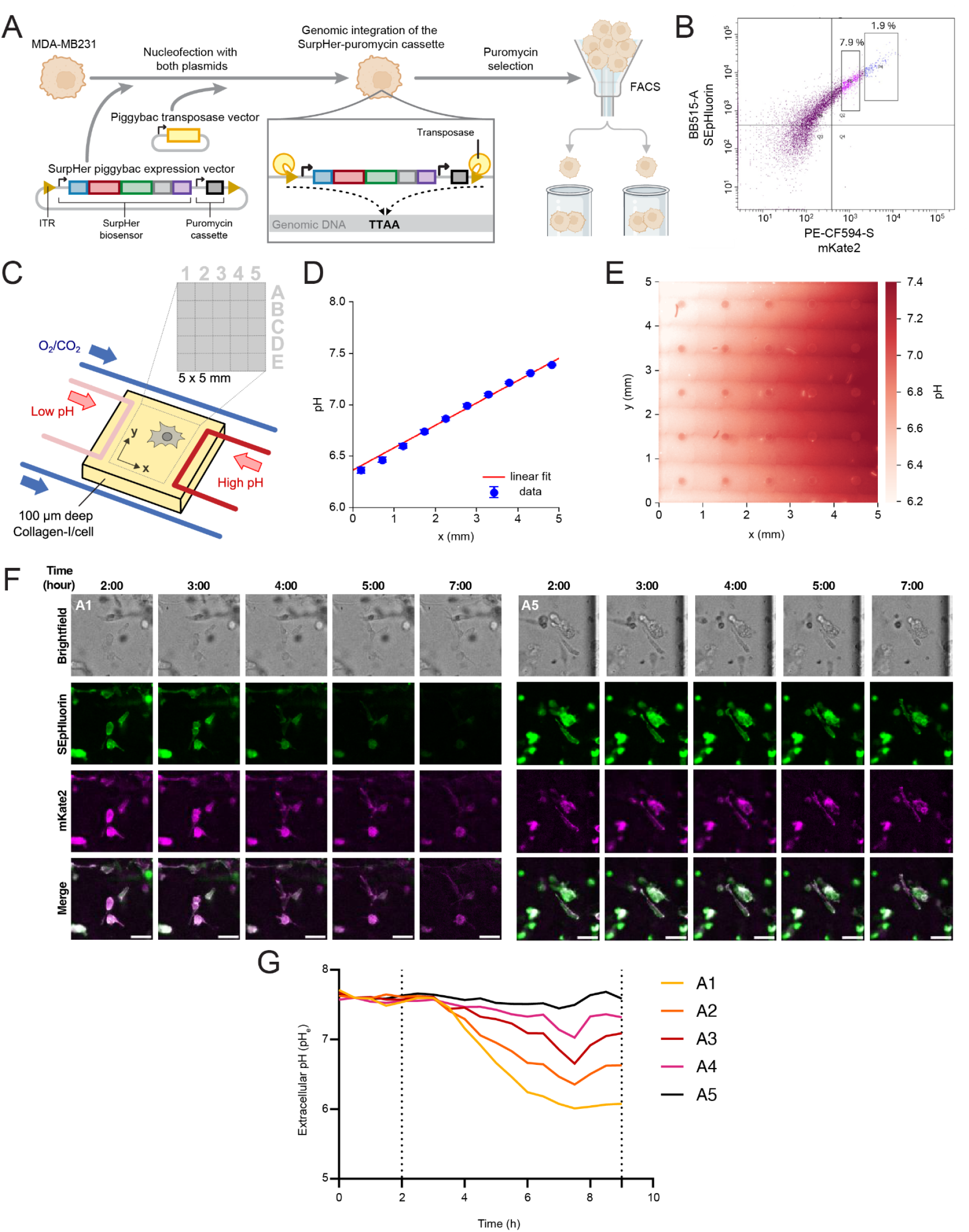
Extracellular pH live imaging in a microfluidic device. **A**. Illustration of cell sorting of transfected MDA-MB-231 cells. **B**. Sorting of the brightest double-positive SEpHluorin+/mKate2+ cells, which represented only around 2% of living cells. **C**. Schematic illustration of liquid and gas channels with regard to the collagen-I/cell slab. The pH gradient establishes along the *x-*direction between the source and drain microchannels. The grid represents the FOVs used for imaging the cells in the observation chamber (5×5 mm^2^). **D**. pH gradient across the observation chamber using Phenol-red-free DMEM medium and SNARF. **E**. Heat map of pH in the observation chamber. **F**. Representative zoom of cells inside the pH gradient in FOVs A1 and A5 over time. **G**. Quantification of pH_e_ through the first row of FOVs over time. 8 to 10 cells analysed per FOVs. The first vertical dotted line indicates the moment the left channel was switched from alkaline to acidic DMEM. Scale bar = 50 µm.

Flow cytometric analysis identified two dual-positive SEpHluorin+/mKate2+ populations: one with low SurpHer biosensor intensity, comprising approximately 8% of the live cells, and a second with higher SurpHer intensity, comprising only 2%. After expansion and validation of the correct plasma membrane localisation of the SurpHer biosensor, the second population with the highest SurpHer biosensor expression was selected for the microfluidic experiment. We next applied the stable MDA-MB-231 SurpHer reporter cell line in a microfluidic platform designed to impose a defined extracellular pH gradient across cells cultured in a collagen film, thereby modelling spatially heterogeneous acidification in the tumour microenvironment (Fig. 3C).

Within the microfluidic device, the pH gradient is established and characterised by ratiometric measurement of SNARF pH indicator across the observation chamber (Fig. 3D-E). The collagen layer containing the stable MDA-MB-231 SurpHer reporter cell line was mounted in the microfluidic chip and perfused with Phenol Red-free DMEM medium adjusted to pH 7.6. Both liquid channels were stably provided with the alkaline medium using pressure-driven pumps for the first 2 hours, after which the medium in the left microchannel was switched to the DMEM of pH 6.0. The fluorescence intensity of the cells in the observation chamber indicates changes and drops in pH. On the acidic side (FOV A1), the SEpHluorin mean fluorescence intensity (MFI) dropped over time, while mKate2 remained mainly stable. On the alkaline side (FOV A5), as expected, the recorded signal in SEpHluorin and mKate2 did not change (Fig. 3F, Supplementary Fig. 10A). A correlated response to the pH change occurs in the rest of the FOVs. See the Supplementary videos which include the first (A1, A5), third (C1, C5), and fifth (E1, E5) rows. To calibrate the SurpHer pH_e_-sensitive change, we perfused reporter cell line-containing collagen slices with Phenol Red-free DMEM media of known pH values (Supplementary Fig. 10B). After gradient stabilisation at around 7 hours, SurpHer fluorescence ratios confirmed the gradient that was consistent with the expected pH_e_ in the observation chamber (Fig. 3G). The spatial distribution of the ratiometric signal, therefore, tracked the direction of the applied pH gradient across the device. Together, these findings showed that stable expression of SurpHer supports extracellular pH imaging in a tumour-relevant microfluidic model and enables visualisation of pH heterogeneity across living cancer cell populations.

## Discussion

Here we present the design, validation, and characterisation of SurpHer, a novel genetically encoded, membrane-anchored ratiometric biosensor for imaging pH_e_. SurpHer combines SEpHluorin with the far-red GFP-derived reference fluorophore mKate2 and a CD59-GPI membrane anchor. SurpHer was designed through a rationally designed modular screen of multiple combinations of SEpHluorin with different red fluorescent reference fluorophores, promoters, leader sequences, linkers and membrane anchors. The resulting proteins were first screened for predominant plasma membrane localisation in HEK293T cells, followed by detailed analysis of pH sensitivity, brightness and photostability across multiple cell types.

The sensor that performed best in localisation and pH responsiveness was the pCMV-CD59 leader-mKate2-SEpHluorin-linker218-CD59 anchor variant - which we named SurpHer. A lesson from this modular optimisation is that it is challenging to predict which exact combination of design elements will work best for a given sensor protein in a given cellular context, and this has to be experimentally tested. The pCMV promoter resulted in less heterogeneous biosensor expression and membrane localisation compared to pEF1a. Specifically, the CD59 leader-CD59 linker combination driven by pCMV resulted in specific membrane localisation of both fluorophores - a key parameter for this sensor. This combination has previously been used for eLACCO1.1 for plasma membrane localisation of a lactate biosensor, showing its plasma membrane specificity in HeLa, HEK293 and T98G cell lines (Nasu *et al*., 2021). Importantly, mKate2-based constructs showed substantially better plasma membrane localisation than mCherry-based constructs, although mCherry tended to provide brighter fluorescence with the filter settings employed and with a lower pH sensitivity (Shaner *et al*., 2004). Consistent with this, intracellular mCherry aggregation has previously been reported by several groups (Katayama *et al*., 2008; Laviv *et al*., 2016).

mKate2 exhibits a high extinction coefficient (ε, 62,500 M^-1^cm^-1^) and quantum yield (φ, 0.4), and thus high molecular brightness (ε·φ/1000, 25), compared to other far red fluorescent proteins. It is photostable, fast-maturing (maturation t_1/2_<20-30 min), and modestly pH sensitive at neutral-to-alkaline pH (pKa 5.4), with peak brightness at alkaline pH values (Pletnev *et al*., 2008; Shcherbo et al., 2009)(Balleza, Kim and Cluzel, 2018). In SurpHer, this modest pH sensitivity was substantially smaller than the SEpHluorin response over the tested extracellular pH range, allowing mKate2 to serve as an effective reference channel.

SEpHluorin is a widely used pH sensitive protein, derived from GFP by the Miesenböck group (Miesenböck, De Angelis and Rothman, 1998; Ng *et al*., 2002). SEpHluorin exhibits an extinction coefficient of 37,000 M^-1^cm^-1^ and a quantum yield of 0.8, and thus a brightness of 30. It is photostable, fast-maturing (maturation t_1/2_ about 15-30 min), and highly pH sensitive at physiological pH (pKa 7) (Balleza, Kim and Cluzel, 2018)(Miesenböck, De Angelis and Rothman, 1998; Ng *et al*., 2002). These properties make SEpHluorin well suited to reporting extracellular pH changes across the physiological and tumour-associated pH range examined in this study.

SurpHer addresses several limitations of the currently available methods for assessing pH_e_, including small molecule dyes such as water-soluble forms of fluorescein or SNARF, membrane-intercalating dyes (Stock *et al*., 2007; Lauritzen *et al*., 2012), and non-ratiometric genetically encoded pH_e_ reporters, and does not require FRET imaging. Furthermore, we show that using the Piggybac system to stably integrate sufficient copies of SurpHer enables single-cell imaging in a microfluidic device under a conventional epifluorescence microscope, without detectable toxicity. This stable-expression strategy avoids repeated exogenous labelling and supports longitudinal imaging formats that are difficult to implement with transiently applied pH probes.

Importantly, the short and similar biomaturation times of SEpHluorin and mKate2 ensure fast and similar turnover rates, minimising problems with fluorophore loss during long-term imaging. The brightness of both fluorophores and the far-red excitation and emission of mKate2 make SurpHer excellently suited for pH_e_ imaging in more complex systems, such as 3D cell culture models, organoids, and *in vivo* conditions, where imaging depth is a critical issue. In vivo, or in co-culture approaches, SurpHer could be expressed selectively in a cell population of interest.

### Limitations of study

The need to genetically modify cells to employ SurpHer obviously carries inherent limitations. Generating stable cell lines is more time-consuming than applying dyes, and the procedure could potentially alter cell phenotype. However, the tightly controlled copy number per cell using the piggybac system should limit such problems in addition to reducing variability of fluorescence intensity. Consistent with this notion, we have not observed detectable effects on cell viability, morphology or other phenotypic traits.

SurpHer measures pHe in the immediate vicinity of the plasma membrane. Our structural model indicates that the extracellular domain has a maximum molecular dimension of approximately 125 Å, representing an upper bound on its extension from the membrane. This is a potential limitation for determining global pHe, however, in the narrow confined spaces of a tissue, this is likely to be a very relevant measure of interstitial pH. An advantage of this localisation is that SurpHer will enable precise measurements of how net acid transport across the cell membrane alters the pericellular pH. Such measurements have been used to monitor roles of specific ion transport proteins in regulating pericellular pH (Stock *et al*., 2007; Lauritzen *et al*., 2012), and to reveal local pH_e_ niches in epithelia (Urra *et al*., 2008). The thickness of the glycocalyx - the dense layer of membrane-bound glycoproteins and glycolipids emanating from the cell surface - varies greatly between cell types, being extremely thick in endothelial cells (up to several μm), generally much thinner in epithelial cells (100-500 nm), and even thinner in leukocytes. Thus, the position of SurpHer within the unstirred layer formed by the pericellular glycocalyx will vary substantially with cell type. This can be both a strength and a shortcoming of SurpHer as a tool, but it will have to be taken into account when interpreting results, as global and pericellular pH can differ substantially in a cell culture (Krähling *et al*., 2009).

In conclusion, we used a rational modular strategy to design and validate a wide range of novel ratiometric pH_e_ sensing tools, resulting in a high-performing sensor, SurpHer, which we showed to be highly plasma membrane specific, bright, with a large dynamic range and good photobleaching properties. Using piggyback technology, SurpHer could be readily expressed at controlled levels across various cell types and was amenable to a range of fluorescent analysis platforms. Our versatile sensor should enable improved understanding of interstitial pH and core mechanisms of pH_e_ regulation, both in model systems and in vivo.

## Materials and Methods

### Cloning of the Biosensor plasmid library

DNA, oligos and primers were purchased from Integrated DNA Technologies. The biosensor plasmid library was constructed by modular Golden Gate assembly, consisting of five parts into a backbone acceptor vector: part 1 (promoter); part 2 (leader sequence); part 3 (fluorophores); part 4 (linker and anchor sequence); and part 5 (polyA signal). Level 0 parts were constructed by amplifying the individual sequences flanked by Esp31 restriction sites, with overhangs that enabled assembly of parts in the following order: promoter-leader-fluorophore-anchor-transcription terminator. Sequences for each Level 0 part and their origin are listed in Supplementary Table 1.

The Golden Gate assembly was carried out by combining all relevant parts in equimolar ratios to the Level 1 acceptor vector (50 ng). The Level 1 acceptor vector was based on the pDisplay-pHoran4 plasmid (Addgene, 61557), in which i) CMV promoter, pHoran4, and polyA signal were replaced by a ccdB cloning cassette flanked by Esp31 sites and ii) Neo/KanR resistance cassette was swapped with PuroR using a homemade Gibson assembly master mixes according to the published protocol (Gibson *et al*., 2009). For Golden Gate assembly, we combined 10x T4 DNA ligase buffer, 100 units of T4 DNA ligase (NEB, M0202M), 5 units of Esp31 (Thermo, FD0454), all the relevant DNA parts 1-5 and Level 1 acceptor vector DNA to a final volume of 10 µl. The reaction was incubated in a thermocycler with the following conditions: 37°C for 5 min; 30 cycles of 37°C for 5 min and 16°C for 10 min; 16°C for 30 min; 37°C for 60 min; 50°C for 5 min; 80°C for 10 min; and then held at 4°C The reactions were transformed into NEB Stable competent E. coli cells (NEB, C3040H). A single colony was picked from each transformation, cultured in 5 mL of LB medium with ampicillin and the plasmid was purified using the NucleoSpin Plasmid Transfection grade kit (Macherey Nagel, 740490.250M). Purified plasmids were quantified using the Nanodrop One (Thermo Fisher) and verified by Sanger sequencing (Eurofins). Sequences for each Level 1 plasmid used in Fig. 1C-F (excluding the Level 1 acceptor vector backbone sequence) are listed in Supplementary Table 2.

### Cell lines and cell culture

MDA-MB-231 cells were a kind gift from Dr. Marie Kveiborg, Copenhagen, Denmark. MDA-MB-231 cells were STR-profile authenticated and tested for contamination by IDEXX BioAnalytics (most recent test, September 2022). PANC-1 cells were obtained from ATCC (CRL-1469) and were STR profile-authenticated and tested for contamination by IDEXX BioAnalytics (latest test, February 2026). HEK293T cells were a kind gift from Dr. James Bryson, Copenhagen, Denmark and tested for contamination by Eurofins (latest test, December 2025). All cell lines were routinely tested and found negative for mycoplasma infection (approximately every 3 months).

MDA-MB-231, PANC-1 and HEK293T cell lines were cultured in DMEM (Thermo, 41966029) supplemented with 10% fetal bovine serum (Sigma, F9665) and 1% Penicillin/Streptomycin (Sigma, P0781). Cells were kept at 37°C in a humidified incubator with 5% CO_2_ and passaged twice a week once confluency reached 70-80%.

### Biosensor library screening in HEK293T cells

HEK293T cells were seeded at a density of 8,000 cells per well in a 96-well optical-bottom black microplate (Thermo, 165305) then transfected 24 hr after with 100 ng of each plasmid using Lipofectamine 3000 (Thermo, L3000008). At 48 h post-transfection, cells were imaged using the ImageXpress Confocal HT.ai (Molecular Devices) with Texas Red, FITC, and transmitted light channels. Images were analysed using a Python script to standardise analysis conditions across all biosensor variants.

### Characterisation of pH sensing dynamics of the biosensor candidate in HEK293T, MDA-MB231 and PANC-1 cells

All three cell lines were seeded at a density of 0.25-0.4 × 10^6^ per dish in WillCo dishes (WillCoWells BV, HBST-3522) coated with 0.01% poly-L-lysine (Sigma, P9155). After 24 h, cells were transfected using the Lipofectamine 3000 kit (Invitrogen, L3000001) according to the manufacturer’s instructions, using 2 µg of plasmid DNA per dish. During transfection, cells were incubated for 5 h in Opti-MEM (Gibco, 31985062) containing the Lipofectamine–DNA–P3000 complex. Subsequently, cells were washed with 1× PBS and incubated in complete growth medium for an additional 24 h prior to imaging.

Ringer solutions contained 130 mM NaCl, 3 mM KCl, 3.3 mM MOPS (Sigma, M1253), 3.3 mM TES (Sigma, T1375) and 5 mM HEPES (Sigma, H3375), 1 mM MgCl2, 0.5 mM CaCl2 and 10 mM NaOH. Prior to imaging, the Ringer solutions were pH adjusted at 37°C to pH 7.8, 7.4, 7.0, 6.4 and 6.0.

Prior to imaging, cells were visually assessed and washed once in 1× PBS, then incubated in the corresponding pH-adjusted Ringer solution. Imaging was performed using a Nikon Ti2 inverted microscope equipped with a Teledyne 01-Kinetix-M-C camera and a stage-top heating unit maintained at 37°C, using a 60x objective. SEpHluorin fluorescence was acquired using 488 nm excitation at 10% laser power and a 540/40 nm emission filter, while mKate2 fluorescence was acquired using 575 nm excitation at 10% laser power and a 698/70 nm emission filter.

Image analysis was performed using Fiji (ImageJ). Cell membranes were manually delineated using the *Freehand Line ROI* tool with a line width of 20 pixels. Fluorescence intensity measurements were obtained, and the SEpHluorin/mKate2 ratio was calculated after background subtraction. The SEpHluorin/mKate2 ratio was normalised to the first measurement point in pH 7.4. Comparison of the linear regressions (Figure 2D) was done using ANCOVA two-tailed test in Prism v.10.

### AlphaFold3 modelling of SurpHer

AlphaFold 3 modelling and theoretical SAXS analysis of the biosensor. The mature extracellular biosensor domain (residues R26-N526) was modelled using AlphaFold 3 (Abramson *et al*., 2024) via the AlphaFold Server (https://alphafoldserver.com/) with default parameters. To assess the effect of including counter-ions on model quality, predictions were carried out under several ionic conditions combining different ratios of Na+ and Cl-ions. For each condition, 10 independent predictions were generated, and model quality was evaluated by the predicted template-modeling (pTM) score. The condition containing 10 Na^+^ and 10 Cl^−^ ions yielded the highest-confidence models, and the prediction with the highest pTM (pTM=0.57) score was selected as the representative structure. Ions were removed prior to all downstream structural analyses.

The radius of gyration (Rg) and the maximum molecular dimension of the ion-free extracellular biosensor domain (excluding the GPI anchor) were calculated using CRYSOL (Svergun, Barberato and Koch, 1995) as implemented in the ATSAS 4.1.2 program suite (Franke, Gräwert and Svergun, 2025). Calculations were performed using default CRYSOL parameters.

For illustrative purposes, the AlphaFold 3 model was combined with a GPI anchor and positioned in the context of a schematic plasma membrane. The GPI anchor was generated using the default core sequence (PI–GlcN–3×Man–EtNP) implemented in the Membrane Builder module of CHARMM-GUI (Jo *et al*., 2008; Wu *et al*., 2014; Lee *et al*., 2019) in the context of a symmetric POPC bilayer (50 POPC molecules per leaflet). The bilayer was not retained in the final figure. The AlphaFold 3-predicted SurpHer biosensor model was positioned manually relative to the GPI anchor in PyMOL (Schrödinger, LLC), and the plasma membrane is depicted schematically. No covalent bond between the protein and the GPI anchor was modelled, and the orientation of the protein with respect to the membrane is illustrative rather than physically optimized. The selected model is included as a Supplemental file.

### Cloning of SurpHer-piggybac plasmid

Primers were designed to amplify the SurpHer biosensor along with a puromycin cassette, flanked by 40 bp of homology to the piggyBac vector in Addgene plasmid 190034. Separately, the piggybac vector was linearised with primers designed to amplify the backbone vector, excluding the Cas13d gene. The SurpHer-puromycin cassette was assembled into the piggybac expression vector using homemade Gibson assembly master mixes according to the published protocol (Gibson *et al*., 2009) then transformed into NEB Stable competent cells (NEB, C3040H) and selected on LB medium containing ampicillin. The plasmid was purified using the NucleoSpin Plasmid purification kit (Macherey Nagel, 740490.250M) and verified using whole-plasmid sequencing (Eurofins, 3094-0ONTC).

### Stable cell line generation and cell sorting

MDA-MB231 cells were electroporated using Lonza SE Cell Line 4D-Nucleofector® X Kit (Lonza, cat. V4XC-1024/V4XC-1012), using a 3:1 vector ratio of SurpHer and PiggyBac transposase (mPB) (Yusa *et al*., 2009) following manufacturer’s instructions. mPB plasmid was a kind gift from Dr. Navneet Vasistha, Copenhagen, Denmark. The cells were puromycin selected using 1 µg/mL for 10 days before cell sorting.

Prior to cell sorting, cells were detached and stained with 1:1000 of blue live/dead (Thermo Fisher, L34961) for 30 min to assess cell viability. mKate+/SEpHluorin+/live-dead-cells were sorted in a 96-well plate (100 cells per well) using a Symphony S6 cell sorter (BD Biosciences) equipped with a 355 nm UV laser (100 mW) coupled with a 410 LP 450/50 nm filter for distinguishing dead cells, a blue 488 nm (200 nW) laser coupled with a 505 LP, 530/30 nm BB515 for visualising SEpHluorin, and a yellow/green 561 nm (200 nW) laser couple with 600 LP, 610/20 nm PE-CF594 PI for visualising mKate. Data was processed using FACSDiva software (BD, version V8.0) and analysed using FlowJo (Version: v10.1). The sorted cells were expanded and used for microfluidic experiments.

### pH sensitivity of SEpHluorin/mKate2 in a microfluidic gradient

Stably transfected MBA-MB231 expressing SurpHer were embedded within 6 mg/mL rat tail collagen-I (Corning, 734-1085) at pH 7.4 with a final concentration of 4,000,000 cells/mL. Collagen with cells was added to the 100 μm-thick frame (7×7 mm^2^) on a positive glass slide (Thermo, K5800AMNZ72). The frame was prepared by cutting a PCR-compatible tape (MicroAmp™ Optical Adhesive Film, ThermoFisher) with a scalpel-cutting machine (Silhouette Cameo) and laminating it onto the glass slide. For gelation, the collagen layer was covered with a thin Fluorinated Ethylene Propylene (FEP) foil and loaded with the weight of a glass slide for 1 h at 37°C. After gelation, the foil was gently removed, and the glass slide with cells embedded in the collagen was kept in DMEM medium for 72 h prior to the microfluidic gradient experiment.

The PDMS microfluidic device is made hydrophylic using oxygen plasma (ATTO, Diener) and reversibly assembled to a polycarbonate (PC) chip holder connected to fluid and gas tubings. Once the fluid tubings are filled with media, the glass slide containing the collagen with embedded cells is aligned on the PDMS device observation chamber. The PC chip holder, the PDMS device, and the glass slide with collagen-embedded cells are secured tightly together with screws on a metal frame designed for the microscope chamber. See the Supplementary note and Supplementary Fig. 9 for the detailed microfluidic workflow.

DMEM media adjusted to pH 6.0 and 7.6 with HCl (1 M, Merck) and NaOH (1 M, VWR) in the incubator (5% CO_2_, 37°C) are introduced into the liquid channels. Temperature during the experiment is maintained at 37°C using a temperature- and CO_2_-controlled environmental chamber (OKOlab, UNO-T-H-PREMIXED). Pressure-based flow controllers (Fluigent, Flow EZ™) were used to control liquid flow. OxyGEN software (Fluigent, version 2.3.2.0) was used to maintain a flow rate of 4 uL/min at about 30 mbar. 20%O_2_/5%CO_2_ was supplied to the gas tubings at 30 mbar.

SurpHer fluorescence intensity ratios were recorded every 30 min for about ten hours using an inverted microscope (Nikon Eclipse Ti2-E, objective 20x/0.70) equipped with a motorised stage. Imaging of the entire observation chamber was achieved using a 5 × 5 grid field of view (FOV) with a size of 2720 × 2720 pixels (907.28 × 907.28 μm). The z-stack in each FOV consists of 25 planes at a 5 μm pitch. SEpHluorin fluorescence was recorded using 488 nm excitation at 10% laser power and a 540/40 nm emission filter, while mKate2 fluorescence was acquired using 575 nm excitation at 10% laser power and a 698/70 nm emission filter. Z-projections, background subtraction, and brightness and contrast adjustments were performed for each FOV using the same parameters. Mean fluorescence intensities of individual cells were analysed over time using TrackMate plug-in (Ershov *et al*., 2022) within Fiji (version: 1.54p). The Cellpose cyto2.0 pre-trained model (Stringer *et al*., 2021) was incorporated to detect and track cells in the mKate2 channel. We analysed tracks that (i) last the whole experiment without frame gap, and (ii) have SEpHluorin/mKate2 ratio >= 1. Acquisition and analysis of the calibration experiment and data were done using the same parameters. The linear regression equation of the calibration data was used to extract the extracellular pH_e_ sensed by cells in the microfluidic chip. For visualisation (Fig. 3F), a single z-slice was chosen, and a script was used to apply the same parameters to the different FOVs across time (reduction of the background, brightness contrast, green and magenta LUTs, and making a merge with the scale bar).

## Data availability

SurpHer plasmids generated in this study will be deposited in Addgene/are available upon request. Plasmid sequences and image analysis scripts are provided in the Supplementary materials. Any additional raw data is available from the corresponding author upon reasonable request.

## Acknowledgements

We gratefully acknowledge N. Vasistha for assistance with piggybac-based construct integration, M. Flinck and T. Larsen for their technical support and early contributions by A. Fuentes and J. Seshadri. We also thank A. E. C. Diaz from the Laboratory Automation Screening and Microscopy Core Facility, University of Copenhagen for assistance with the high-throughput confocal imaging using the ImageXpress Confocal HT.ai and the Flow Cytometry & Single Cell Core Facility, University of Copenhagen for their assistance with single cell sorting.

## Author contributions

S.C.H., L.D.S., J.Y.A. and S.F.P. conceived the study. S.C.H., J.Y.A., L.D.S., R.C. designed and performed the biosensor screen and characterisation experiments. L.D.S. and M.R.C performed the microfluidics experiments. O.N.E. and B.B.K. performed the AlphaFold3 modelling. S.C.H., J.Y.A., L.D.S., M.R.C., R.M., A.S. and S.F.P. analysed the data and contributed to data interpretation. S.C.H., J.Y.A., M.R.C., R.M., and L.D.S. prepared the figures and wrote the first draft of the manuscript. S.C.H., L.D.S., M.R.C., O.N.E., B.B.K., R.M., A.S., J.Y.A. and S.F.P. reviewed and edited the manuscript. All authors read and approved the final manuscript.

## References

Abramson, J. et al. (2024) ‘Accurate structure prediction of biomolecular interactions with AlphaFold 3’, Nature, 630(8016), pp. 493–500.

Anemone, A. et al. (2019) ‘Imaging tumor acidosis: a survey of the available techniques for mapping in vivo tumor pH’, Cancer metastasis reviews, 38(1-2), pp. 25–49.

Balleza, E., Kim, J.M. and Cluzel, P. (2018) ‘Systematic characterization of maturation time of fluorescent proteins in living cells’, Nature methods, 15(1), pp. 47–51.

Corbet, C. et al. (2020) ‘TGFβ2-induced formation of lipid droplets supports acidosis-driven EMT and the metastatic spreading of cancer cells’, Nature communications, 11(1), p. 454.

Crouigneau, R. et al. (2024) ‘Mimicking and analyzing the tumor microenvironment’, Cell reports methods, 4(10), p. 100866.

Czaplinska, D. et al. (2023) ‘Crosstalk between tumor acidosis, p53 and extracellular matrix regulates pancreatic cancer aggressiveness’, International journal of cancer, 152(6), pp. 1210–1225.

Ershov, D. et al. (2022) ‘TrackMate 7: integrating state-of-the-art segmentation algorithms into tracking pipelines’, Nature methods, 19(7), pp. 829–832.

Franke, D., Gräwert, T. and Svergun, D.I. (2025) ‘New features in, a program suite for small-angle scattering data analysis’, Journal of applied crystallography, 58(Pt 3), pp. 1027–1033.

Gibson, D.G. et al. (2009) ‘Enzymatic assembly of DNA molecules up to several hundred kilobases’, Nature methods, 6(5), pp. 343–345.

Jensen, L.J. et al. (1997) ‘Proton pump activity of mitochondria-rich cells. The interpretation of external proton-concentration gradients’, The Journal of general physiology, 109(1), pp. 73–91.

Jo, S. et al. (2008) ‘CHARMM-GUI: a web-based graphical user interface for CHARMM’, Journal of computational chemistry, 29(11), pp. 1859–1865.

Karsten, L. et al. (2022) ‘Genetically encoded ratiometric pH sensors for the measurement of intra- and extracellular pH and internalization rates’, Biosensors, 12(5), p. 271.

Katayama, H. et al. (2008) ‘GFP-like proteins stably accumulate in lysosomes’, Cell structure and function, 33(1), pp. 1–12.

Ke, G. et al. (2014) ‘A cell-surface-anchored ratiometric fluorescent probe for extracellular pH sensing’, ACS Applied Materials & Interfaces, 6(17), pp. 15329–15334.

Krähling, H. et al. (2009) ‘The glycocalyx maintains a cell surface pH nanoenvironment crucial for integrin-mediated migration of human melanoma cells’, Pflugers Archiv : European journal of physiology, 458(6), pp. 1069–1083.

Kuner, T. and Augustine, G.J. (2000) ‘A genetically encoded ratiometric indicator for chloride: capturing chloride transients in cultured hippocampal neurons’, Neuron, 27(3), pp. 447–459.

Lauritzen, G. et al. (2012) ‘The Na+/H+ exchanger NHE1, but not the Na+, HCO3(-) cotransporter NBCn1, regulates motility of MCF7 breast cancer cells expressing constitutively active ErbB2’, Cancer letters, 317(2), pp. 172–183.

Laviv, T. et al. (2016) ‘Simultaneous dual-color fluorescence lifetime imaging with novel red-shifted fluorescent proteins’, Nature methods, 13(12), pp. 989–992.

Lee, J. et al. (2019) ‘CHARMM-GUI Membrane Builder for Complex Biological Membrane Simulations with Glycolipids and Lipoglycans’, Journal of chemical theory and computation, 15(1), pp. 775–786.

Miesenböck, G., De Angelis, D.A. and Rothman, J.E. (1998) ‘Visualizing secretion and synaptic transmission with pH-sensitive green fluorescent proteins’, Nature, 394(6689), pp. 192–195.

Nasu, Y. et al. (2021) ‘A genetically encoded fluorescent biosensor for extracellular L-lactate’, Nature communications, 12(1), p. 7058.

Ng, M. et al. (2002) ‘Transmission of olfactory information between three populations of neurons in the antennal lobe of the fly’, Neuron, 36(3), pp. 463–474.

Pletnev, S. et al. (2008) ‘A crystallographic study of bright far-red fluorescent protein mKate reveals pH-induced cis-trans isomerization of the chromophore’, The Journal of biological chemistry, 283(43), pp. 28980–28987.

Shaner, N.C. et al. (2004) ‘Improved monomeric red, orange and yellow fluorescent proteins derived from Discosoma sp. red fluorescent protein’, Nature biotechnology, 22(12), pp. 1567–1572.

Shcherbo, D. et al. (2009) ‘Far-red fluorescent tags for protein imaging in living tissues’, The Biochemical journal, 418(3), pp. 567–574.

Shen, Y. et al. (2014) ‘pHuji, a pH-sensitive red fluorescent protein for imaging of exo- and endocytosis’, The Journal of Cell Biology, 207(3), pp. 419–432.

Stigliani, A. et al. (2024) ‘Adaptation to an acid microenvironment promotes pancreatic cancer organoid growth and drug resistance’, Cell reports, 43(7), p. 114409.

Stock, C. et al. (2007) ‘pH nanoenvironment at the surface of single melanoma cells’, Cellular physiology and biochemistry : international journal of experimental cellular physiology, biochemistry, and pharmacology, 20(5), pp. 679–686.

Stringer, C. et al. (2021) ‘Cellpose: a generalist algorithm for cellular segmentation’, Nature methods, 18(1), pp. 100–106.

Svergun, D., Barberato, C. and Koch, M.H.J. (1995) ‘CRYSOL - A program to evaluate x-ray solution scattering of biological macromolecules from atomic coordinates’, Journal of Applied Crystallography, 28(6), pp. 768–773.

Swietach, P., Boedtkjer, E. and Pedersen, S.F. (2023) ‘How protons pave the way to aggressive cancers’, Nature Reviews Cancer, 23(12), pp. 825–841.

Urra, J. et al. (2008) ‘A genetically encoded ratiometric sensor to measure extracellular pH in microdomains bounded by basolateral membranes of epithelial cells’, Pflugers Archiv : European journal of physiology, 457(1), pp. 233–242.

Whitlow, M. et al. (1993) ‘An improved linker for single-chain Fv with reduced aggregation and enhanced proteolytic stability’, Protein engineering, 6(8), pp. 989–995.

Wu, E.L. et al. (2014) ‘CHARMM-GUI Membrane Builder toward realistic biological membrane simulations’, Journal of computational chemistry, 35(27), pp. 1997–2004.

Yang, L. et al. (2021) ‘High-Throughput and Real-Time Monitoring of Single-Cell Extracellular pH Based on Polyaniline Microarrays’, Analytical chemistry, 93(41), pp. 13852–13860.

Yang, Y. et al. (2018) ‘A cell-surface-specific ratiometric fluorescent probe for extracellular pH sensing with solid-state fluorophore’, ACS Sensors, 3(11), pp. 2278–2285.

Yusa, K. et al. (2009) ‘Generation of transgene-free induced pluripotent mouse stem cells by the piggyBac transposon’, Nature methods, 6(5), pp. 363–369.

