## Supplementary Materials for "SurpHer: a genetically encoded ratiometric sensor for dynamic extracellular pH imaging"

**List of supplementary materials**

Supplementary Figures 1-8

Supplementary note: microfluidic device workflow

Supplementary Figures 9-10

Supplementary tables

Supplementary videos

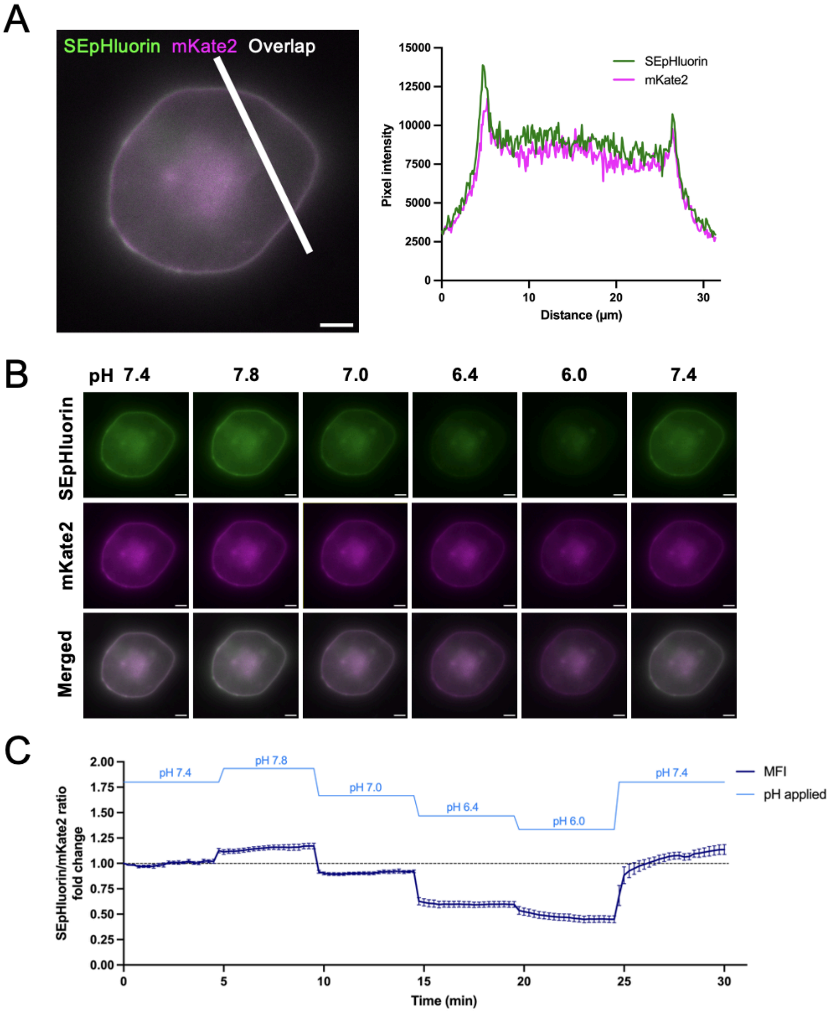


**Supplementary Figure 1 – Variant 1 pH$_e$-sensitive genetically encoded ratiometric biosensor validation in 2D HEK293-T cells. A)** Representative validation of membrane localization of the biosensor. SEpHluorin in green, mKate2 in magenta and overlap in white. Scale bar: 5 µm. The white line represent the ROI for the profile plot analysis. **B)** Representative images of transiently transfected HEK293-T cells in pH 7.8, 7.4, 7.0, 6.4 and 6.0. Imaged with 60x objective. pH insensitive fluorophore: mKate2 (ex: 588 nm, em: 633 nm). pH sensitive fluorophore: SEpHluorin (ex: 488 nm, em: 511 nm). Scale bar: 5 µm. **C)** Quantification of pHe-sensitivity normalized to first measurement in pH 7.4. Images acquired every 15 sec for a total of 30 min. Data analysis by ROI marking of the cell membrane for intensity measurement during pH exposures over time. n = 3, SEM error bars.

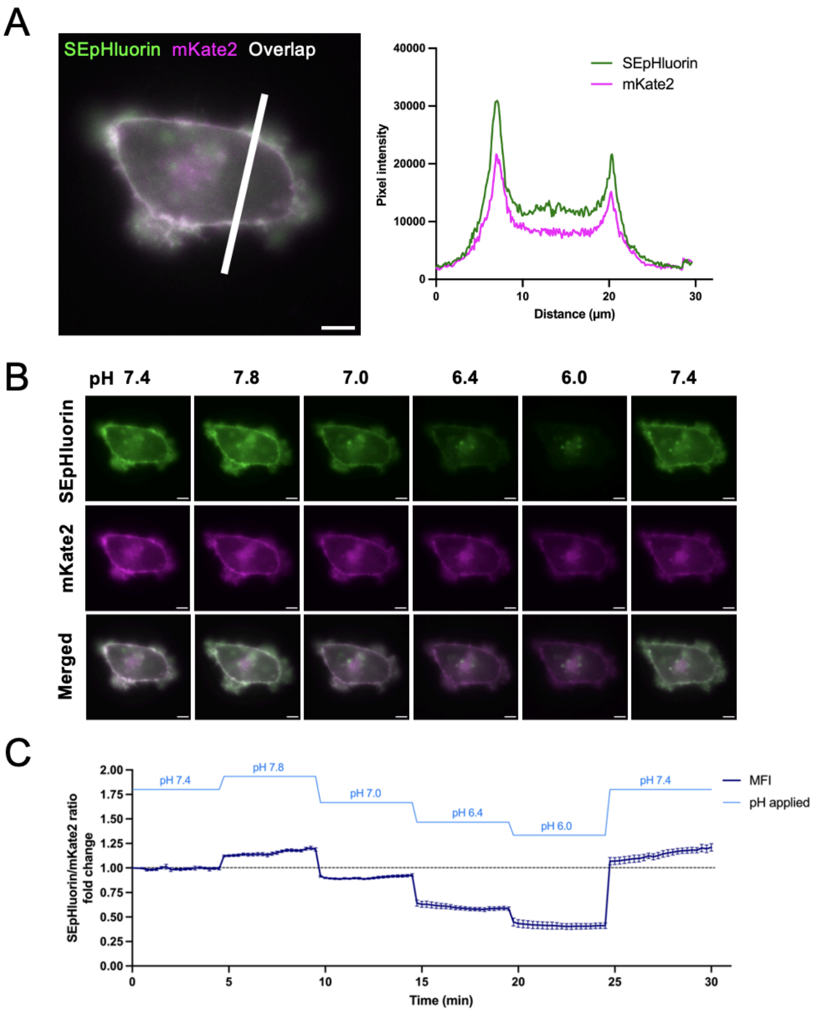


**Supplementary Figure 2 – Variant 1 pH$_e$-sensitive genetically encoded ratiometric biosensor validation in 2D MDA-MB-231 cells. A)** Representative validation of membrane localization of the biosensor. SEpHluorin in green, mKate2 in magenta and overlap in white. Scale bar: 5 µm. The white line represent the ROI for the profile plot analysis. **B)** Representative images of transiently transfected MDA-MB-231 cells in pH 7.8, 7.4, 7.0, 6.4 and 6.0. Imaged with 60x objective. pH insensitive fluorophore: mKate2 (ex: 588 nm, em: 633 nm). pH sensitive fluorophore: SEpHluorin (ex: 488 nm, em: 511 nm). Scale bar: 5 µm. **C)** Quantification of pHe-sensitivity normalized to first measurement in pH 7.4. Images acquired every 15 sec for a total of 30 min. Data analysis by ROI marking of the cell membrane for intensity measurement during pH exposures over time. n = 3, SEM error bars.

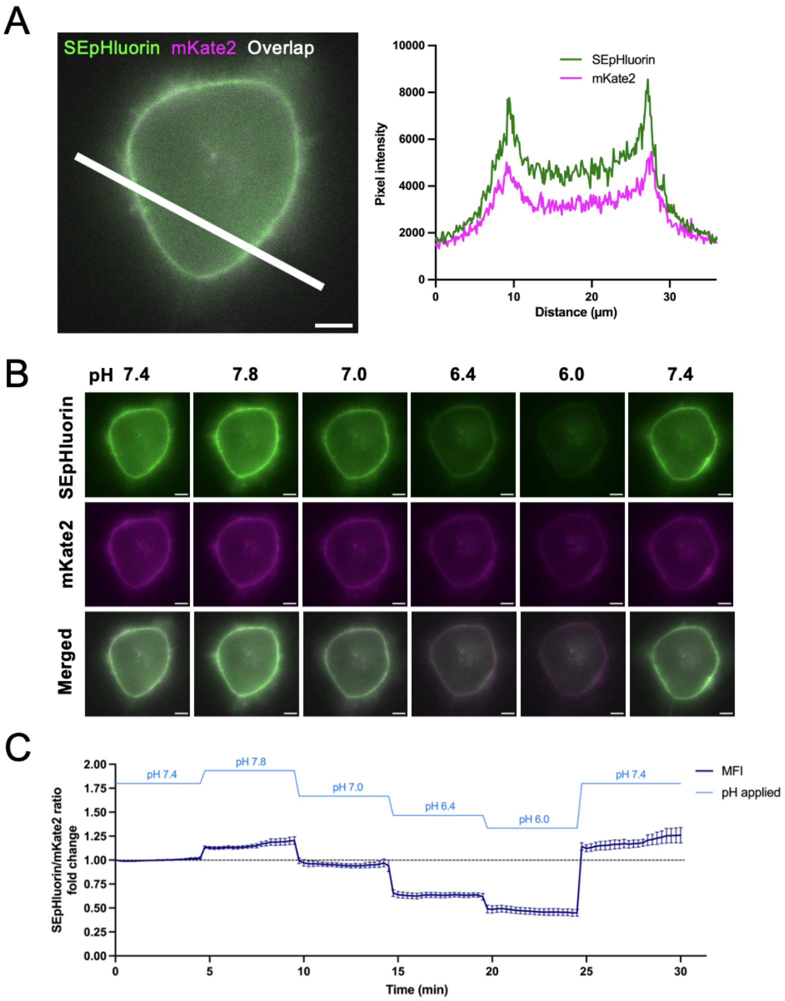


**Supplementary Figure 3 - Variant 1 pH$_e$-sensitive genetically encoded ratiometric biosensor validation in 2D PANC-1 cells. A)** Representative validation n of membrane localization of the biosensor. SEpHluorin in green, mKate2 in magenta and overlap in white. Scale bar: 5 µm. The white line represent the ROI for the profile plot analysis. **B)** Representative images of transiently transfected PANC-1 cells in pH 7.8, 7.4, 7.0, 6.4 and 6.0. Imaged with 60x objective. pH insensitive fluorophore: mKate2 (ex: 588 nm, em: 633 nm). pH sensitive fluorophore: SEpHluorin (ex: 488 nm, em: 511 nm). Scale bar: 5 µm. **C)** Quantification of pHe-sensitivity normalized to first measurement in pH 7.4. Images acquired every 15 sec for a total of 30 min. Data analysis by ROI marking of the cell membrane for intensity measurement during pH exposures over time. n = 3, SEM error bars.

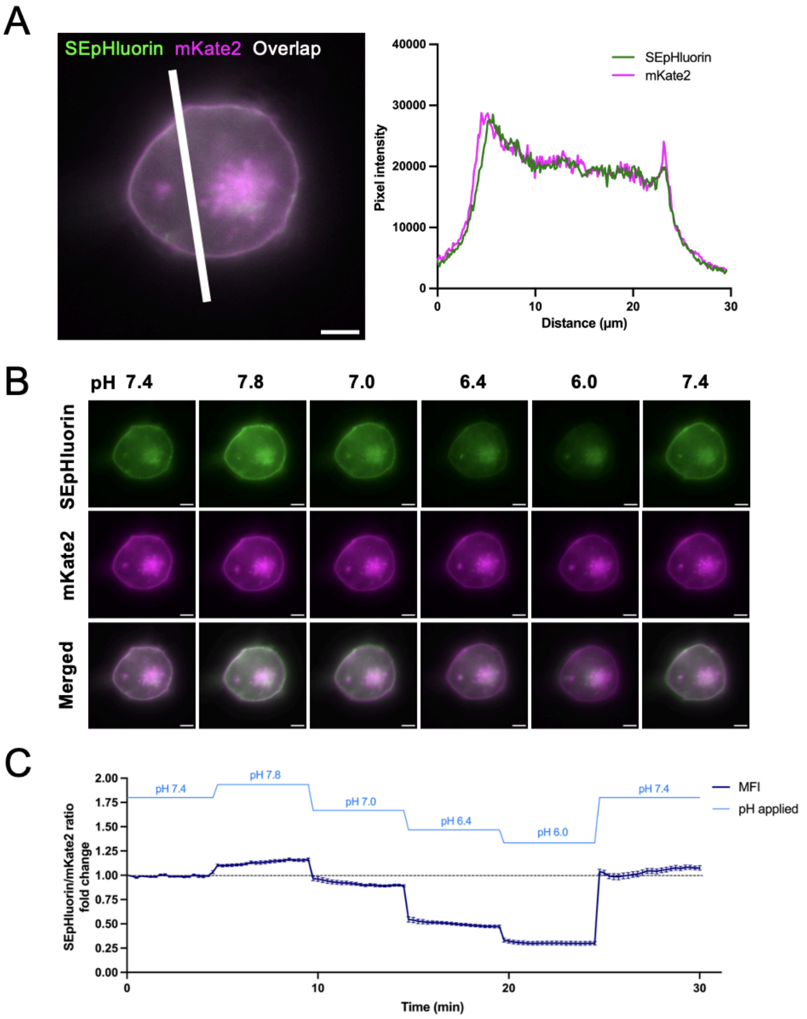


**Supplementary Figure 4 - Variant 2 $pH_e$-sensitive genetically encoded ratiometric biosensor validation in 2D HEK293-T cells. A)** Representative validation of membrane localization of the biosensor. SEpHluorin in green, mKate2 in magenta and overlap in white. Scale bar: 5 µm. The white line represent the ROI for the profile plot analysis. **B)** Representative images of transiently transfected HEK293-T cells in pH 7.8, 7.4, 7.0, 6.4 and 6.0. Imaged with 60x objective. pH insensitive fluorophore: mKate2 (ex: 588 nm, em: 633 nm). pH sensitive fluorophore: SEpHluorin (ex: 488 nm, em: 511 nm). Scale bar: 5 µm. **C)** Quantification of pHe-sensitivity normalized to first measurement in pH 7.4. Images acquired every 15 sec for a total of 30 min. Data analysis by ROI marking of the cell membrane for intensity measurement during pH exposures over time. n = 3, SEM error bars.

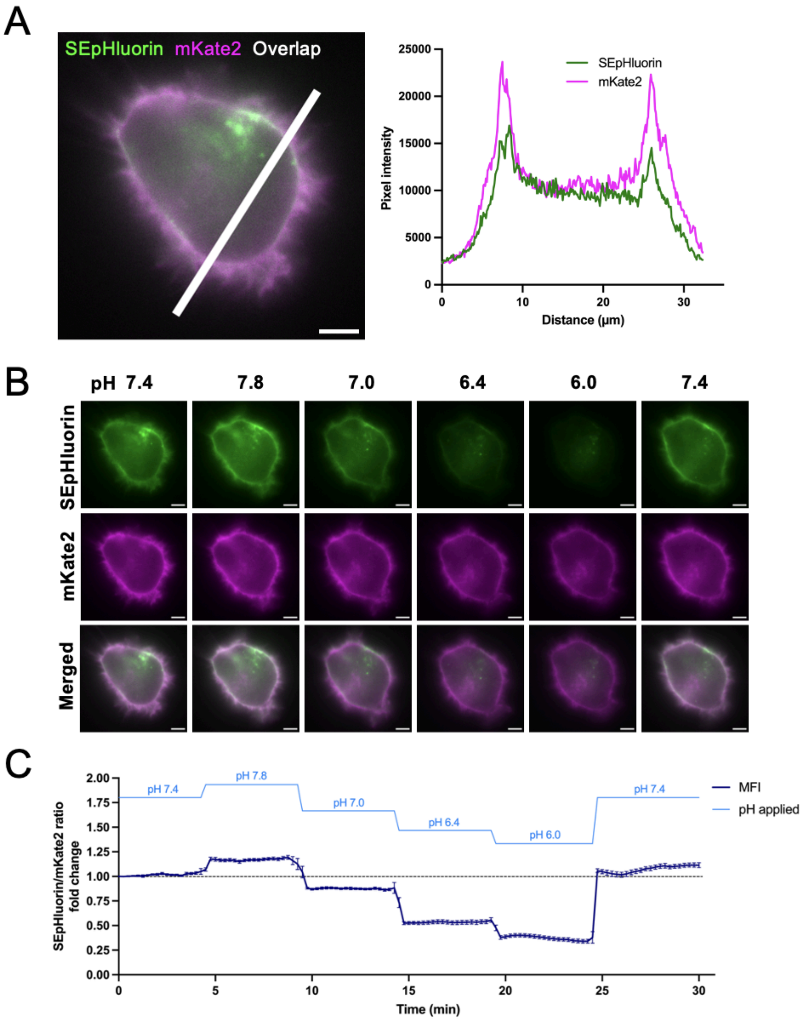


**Supplementary Figure 5 – Variant 2 pH$_e$-sensitive genetically encoded ratiometric biosensor validation in 2D PANC-1 cells. A)** Representative validation of membrane localization of the biosensor. SEpHluorin in green, mKate2 in magenta and overlap in white. Scale bar: 5 µm. The white line represent the ROI for the profile plot analysis. **B)** Representative images of transiently transfected PANC-1 cells in pH 7.8, 7.4, 7.0, 6.4 and 6.0. Imaged with 60x objective. pH insensitive fluorophore: mKate2 (ex: 588 nm, em: 633 nm). pH sensitive fluorophore: SEpHluorin (ex: 488 nm, em: 511 nm). Scale bar: 5 µm. **C)** Quantification of pHe-sensitivity normalized to first measurement in pH 7.4. Images acquired every 15 sec for a total of 30 min. Data analysis by ROI marking of the cell membrane for intensity measurement during pH exposures over time. n = 3, SEM error bars.

A
B
C

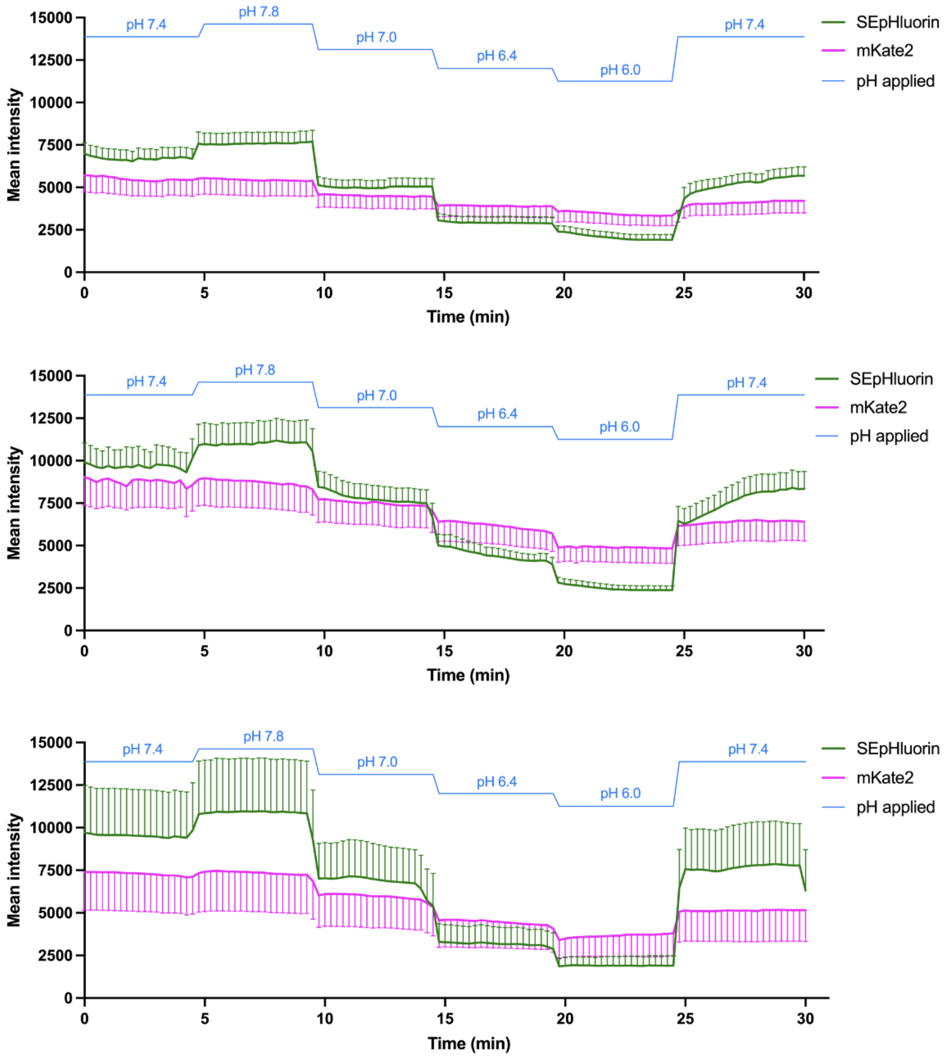


**Supplementary Figure 6 – Variant 1 pHe-sensitive genetically encoded ratiometric biosensor validation in 2D HEK293-T, MDA-MB-231 and PANC-1 cells.** Quantification of pHe-sensitivity of the individual fluorophores in (A) HEK293-T cells, (B) MDA-MB-231 cells and (C) PANC-1 cells, n = 3, SEM error bars. Images acquired every 15 sec for a total of 30 min. Data analysis by ROI marking of the cell membrane for intensity measurement during pH exposures over time. Individual channels shown: SEpHluorin in green and mKate2 in magenta. pH exposure shown in blue with the corresponding pH values annotated above.

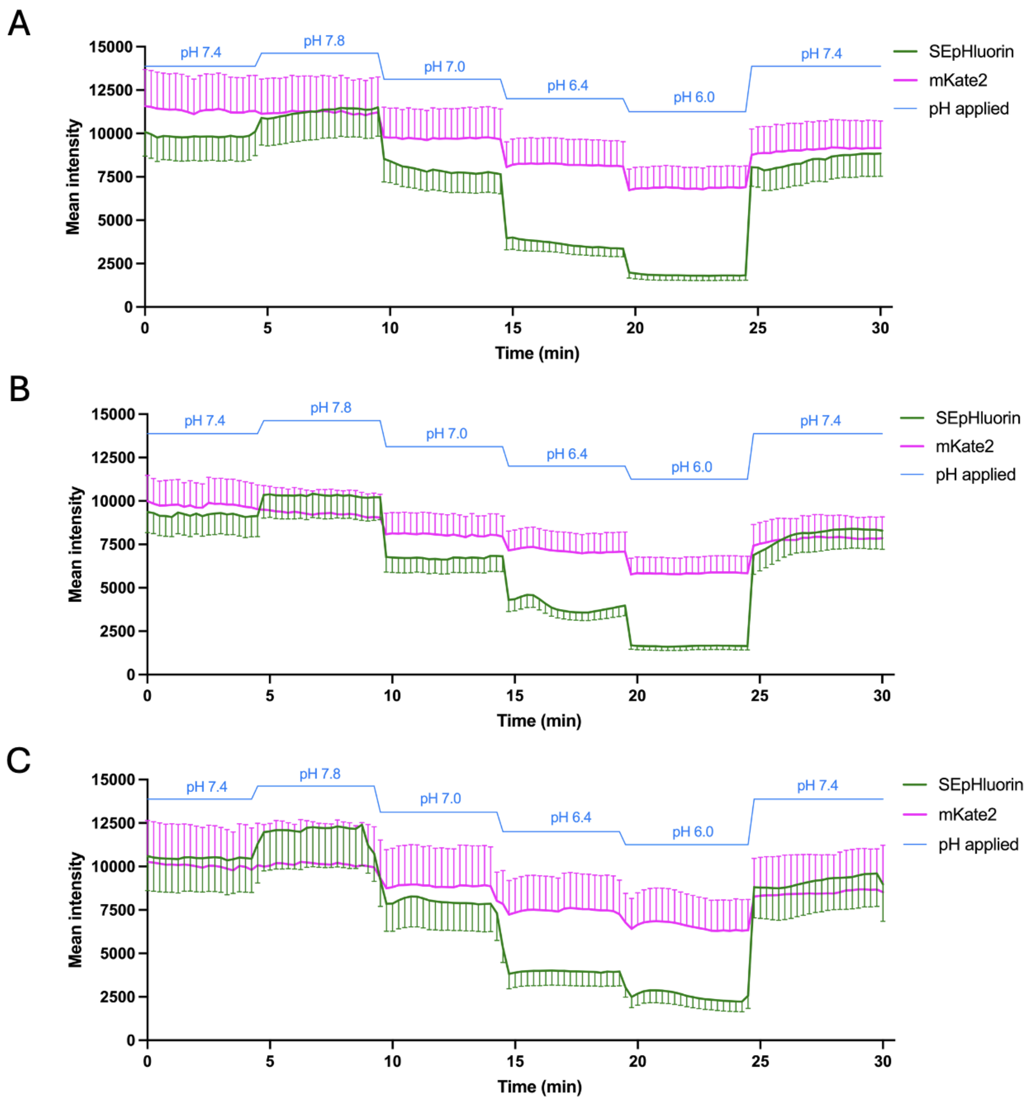


**Supplementary Figure 7 – Variant 2 / SurpHer pH$_e$-sensitive genetically encoded ratiometric biosensor validation in 2D HEK293-T, MDA-MB-231 and PANC-1 cells.** Quantification of pH$_e$-sensitivity of the individual fluorophores in (A) HEK293-T cells, (B) MDA-MB-231 cells and (C) PANC-1 cells, n = 3, SEM error bars. Images acquired every 15 sec for a total of 30 min. Data analysis by ROI marking of the cell membrane for intensity measurement during pH exposures over time. Individual channels shown: SEpHluorin in green and mKate2 in magenta. pH exposure shown in blue with the corresponding pH values annotated above.

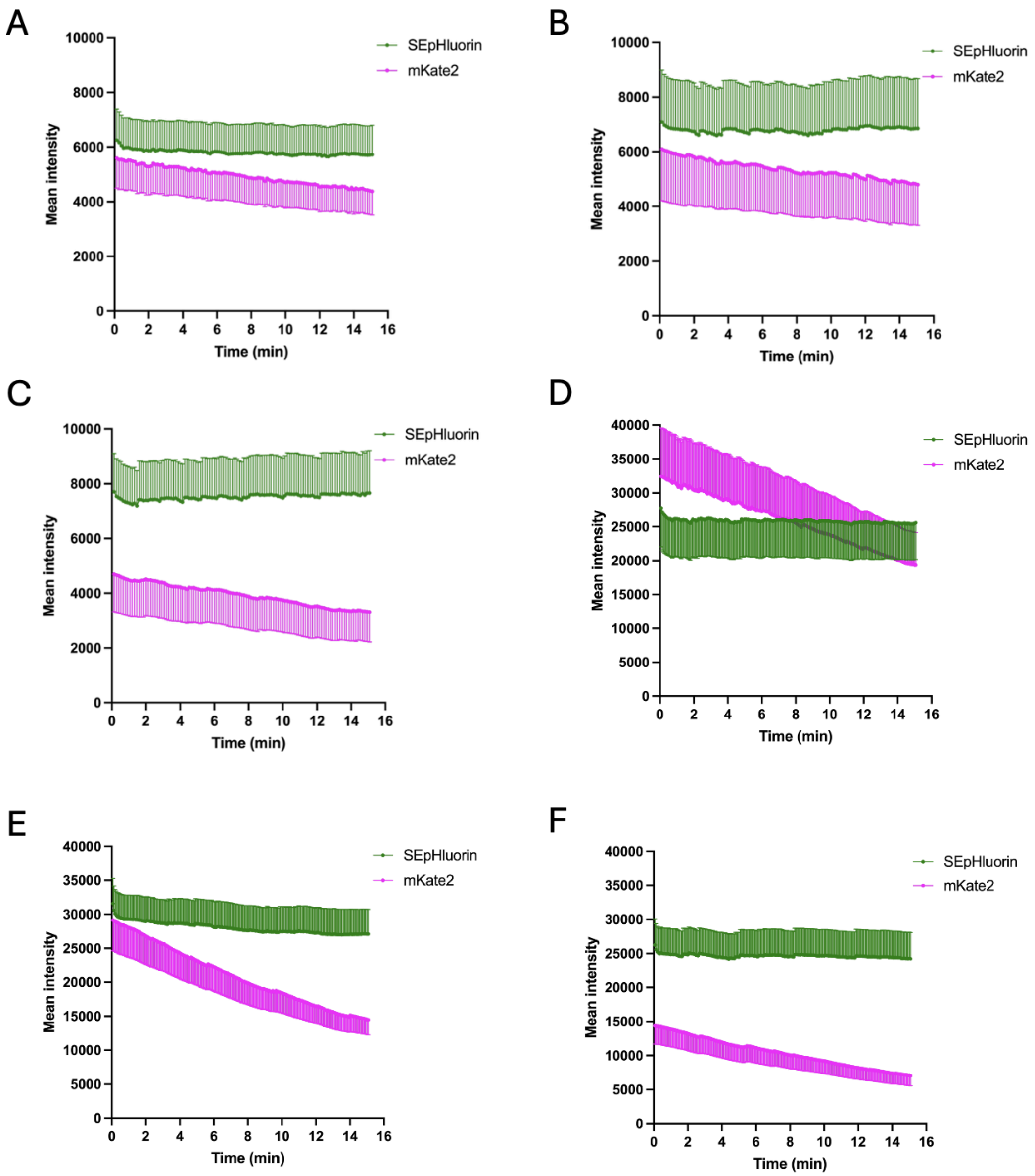


**Supplementary Figure 8 - Normal-intensity and high-intensity bleaching test of the Variant 2 / SurpHer $pH_e$-sensitive genetically encoded ratiometric biosensor in 2D HEK293-T, MDA-MB-231 and PANC-1 cells.** Quantification of bleaching response of the individual fluorophores. Normal-intensity bleaching test in **(A)** HEK293-T cells, **(B)** MDA-MB-231 cells and **(C)** PANC-1 cells. Data analysis by ROI marking of the cell membrane for intensity measurement during normal-intensity bleaching: exposure time corresponding to ~3200 mean intensity with 10% laser exposure for both SEpHluorin and mKate2, imaging every $8^{th}$ second for 15 minutes in pH 7.4. n = 2, SEM error bars. High-intensity bleaching test in **(D)** HEK293-T cells, **(E)** MDA-MB-231 cells and **(F)** PANC-1 cells: exposure time corresponding to ~10.000 mean intensity with 10% laser exposure for both SEpHluorin (150 ms) and mKate2 (300 ms), imaging every $4^{th}$ second for 15 minutes in pH 7.4. Individual channels shown: SEpHluorin in green and mKate2 in magenta. n =2, SEM error bars.

**Supplementary note:**

*PDMS device fabrication*: The Poly(dimethylsiloxane) (PDMS) microfluidic device consists of a single layer (Supplementary Fig. 9A) fabricated by casting PDMS onto a photresistmould. Briefly, the mould was prepared using SU-8 photolithography on a silicon substrate, creating a 100 μm-thick SU-8 structure on a thin SU-8 layer. PDMS was prepared using a SYLGARD™ 184 Silicone Elastomer Kit, with the monomer and crosslinker mixed at a weight ratio of 10:1. The mixture was degassed, then poured onto the mould and cured at 80°C for 2 h. After curing, the PDMS slab was peeled off the mould and cut to the size of a glass slide (76 mm x 26 mm). Fluidic connections to the liquid flow channels and gas connections to the gas flow channels were made by punching holes through the PDMS using a tissue biopsy punch (Harris Uni-core, diameter 1.20 mm for liquid connections and 2 mm for gas connections).

A 5x5 $mm^2$ observation chamber covers the gradient area, two liquid channels, and two gas flow channels, symmetrically positioned relative to the gradient observation area (Supplementary Fig. 9A). The liquid channels are 100 μm wide and 100 μm deep, separated from the gradient observation area by a 500 μm gap. The gas channels in the plane of the gradient observation area are positioned 2 mm away from it to prevent overlap with the collagen layer, thereby increasing the robustness of the assembly (Supplementary Fig. 9B). The gas and liquid channels, as well as the gradient observation area, are separated by PDMS.

*Device assembly and mounting*: The PDMS device and the glass slide with collagen were held together by a homemade Poly(carbonate) (PC) holder that comprised fluidic connectors and a metal frame. On the glass slide, a PCR-compatible tape (100 μm thick) is laminated to accommodate the collagen layer in a 7x7 $mm^2$ cut area. Four 3-mm screws are tightened to a maximum torque of 4 cN·m to maintain reversible sealing. Pressure-based flow controllers (Fluigent, Flow EZ™) were used to drive the liquid flow. The pH of the solutions (6.0 and 7.6) was adjusted by adding the required amounts of HCl and NaOH to the cell culture media (DMEM, 41966-029, Gibco) and was subsequently equilibrated in the incubator (37°C, 5% $CO_2$, 21% $O_2$). The tubing connecting the flow controllers, flow rate sensors, and the device holder were pre-filled with solutions. Next, the oxygen plasma-treated PDMS device was placed on the top of the PC holder with all inlet and outlet holes aligned. The PDMS microfluidic device was then covered with cell culture medium before the glass slide, with the collagen-I film facing down, was assembled on top of the PDMS to avoid trapped air. The collagen-I layer was aligned with the gradient observation area of the PDMS device and with the gaps between the gas and liquid flow channels (Supplementary Fig. 9C). Finally, the gas channels were connected to an $O_2/CO_2$ supply. A gas mixture of 20% oxygen and 5% $CO_2$ is supplied to ensure the necessary conditions for cell growth and the establishment of a $CO_2$-buffered pH, in addition to the microscope enclosure gas control.

*pH gradient and calibration curve*: The pH gradient in the observation chamber was visualized and quantified in a separate experiment using 5 µM of 5-(and-6)-Carboxy SNARF-1 (#C1270, ThermoFisher) in DMEM cell culture medium. Ratiometric measurement was performed using excitation at 488 nm and emission filters of 582/75 nm and 698/70 nm (Supplementary Fig. 9D). The ratio was measured along the *x*-direction of the observation area.

To determine the pH values across the observation area, a calibration curve was generated in yet a separate experiment (Supplementary Fig. 9E). The SNARF ratiometric signal was determined as a function of pH, using four pH-adjusted DMEM media (Phenol-red free). In this calibration experiment, separate collagen slides were soaked overnight at their respective pH with SNARF dye before being mounted in the device. For each collagen slide, the ratiometric signal was recorded after the following conditions were met: (i) respective pH-adjusted medium flowing in both liquid channels, (ii) connected gas channels and $CO_2$ supply of the enclosure, and (iii) temperature at 37°C.

We also investigated the gradient across the observation chamber using Ringer solution (115 mM NaCl, 5 mM KCl, 1 mM $Na_2HPO_4$, 1 mM $CaCl_2$, 0.5 mM $MgCl_2$, and 24 mM $NaHCO_3$). The ratiometric measurements are shown in Supplementary Figs. 9G-H.

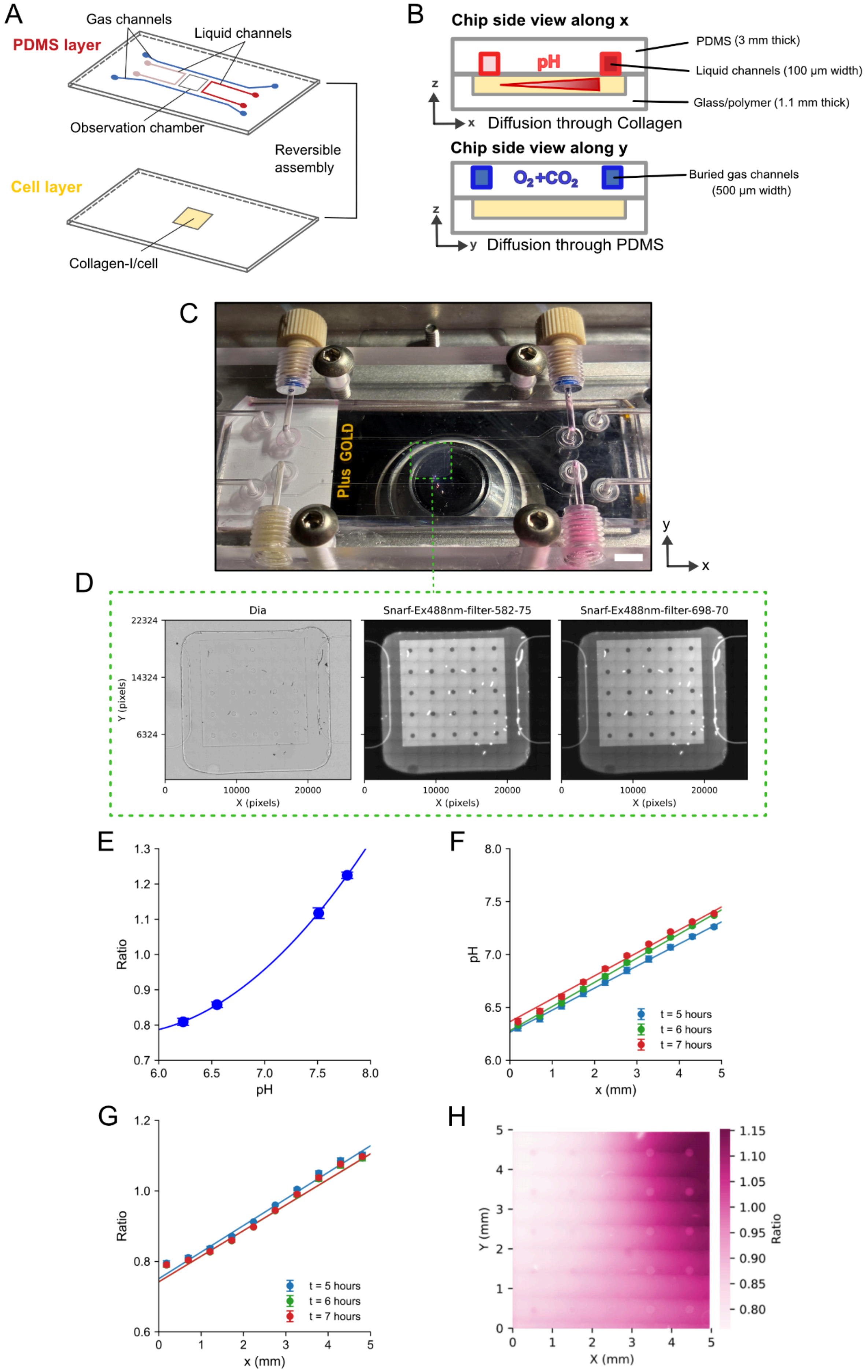
A
Gas channels
Liquid channels
PDMS layer
Observation chamber
Reversible assembly
Cell layer
Collagen-I/cell
B
Chip side view along x
PDMS (3 mm thick)
pH
Liquid channels (100 µm width)
Glass/polymer (1.1 mm thick)
Diffusion through Collagen
Chip side view along y
O2+CO2
Buried gas channels (500 µm width)
Diffusion through PDMS
C
Plus GOLD
D
Dia
Snarf-Ex488nm-filter-582-75
Snarf-Ex488nm-filter-698-70
Y (pixels)
X (pixels)
E
Ratio
pH
F
pH
x (mm)
t = 5 hours
t = 6 hours
t = 7 hours
G
Ratio
x (mm)
t = 5 hours
t = 6 hours
t = 7 hours
H
Y (mm)
X (mm)
Ratio

**Supplementary Figure 9 - Microfluidic pH gradient characterization.** (A) Schematics of the PDMS layer design and reversible assembly to the glass slide hosting the collagen/cell layer. (B) Side views of the assembled PDMS device, including the diffusion paths of the liquid and gas. (C) Picture of the device mounted using the PC holder and metal frame, and placed under the microscope. Scale bar is 5 mm. (D) Large bright field and fluorescent (emission at 582/75 nm and 698/70 nm) images of the collagen area with 20X objective. (E) Calibration of the SNARF ratiometric signal in DMEM at 37 °C in the presence of 5% $CO_2$. (F) pH measured along direction *x* in the observation area at t = 5, 6, and 7 hours after the onset of liquid and gas flow. (G) SNARF ratiometric signal along direction *x* in the observation area at t = 5, 6, and 7 hours after the onset of liquid using Ringer solution. (H) Heat map of the data in (G).

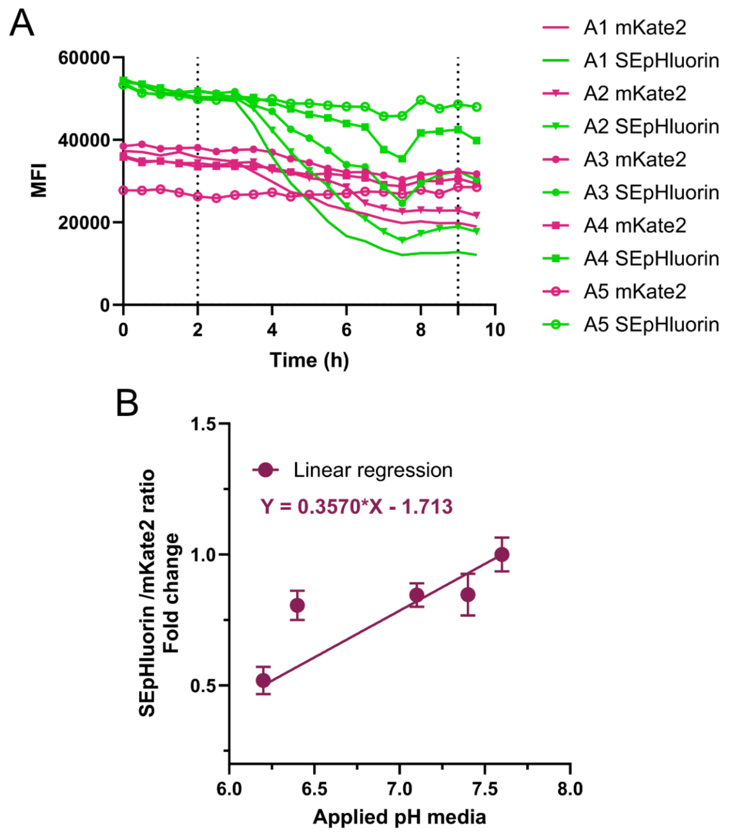


**Supplementary Figure 10 - SurpHer calibration in collagen for microfluidic**. **A.** Mean fluorescence intensity of SEpHluorin and mKate2 over time across the first row of FOVs. 8 to 10 cells were analysed per FOV. **B.** SEpHluorin/mKate2 fold change upon known applied pH media. DMEM Phenol-red free media adjusted and equilibrated in the incubator overnight at pH 7.6, 7.4, 7.2, 6.4 and 6.2. Three hours before image acquisition, media were incubated on several collagen slices containing MDA-MB-231 stably transfected with SurpHer. Collagen slices were then imaged one by one inside the microscope chamber, where temperature (37°C) and 5% $CO_2$ supply were stable (15min). Every cell's SEpHluorin/mKate2 ratio is then normalised to the pH 7.6 mean ratio. The purple line represents the linear regression, ending at pH 6.2.  3 field of views (FOVs) were analysed with a total of 30 to 55 cells per applied pH. Error bars indicate the standard deviation (SD).
